# Interference with MHC class I epitope trimming provides paradoxical protection from autoimmune diabetes

**DOI:** 10.64898/2026.02.19.706793

**Authors:** Barbara Bertocci, Emmanuelle Waeckel-Énée, Nadia Keelan, Sylvaine You, Pierre David, Peter van Endert

## Abstract

Polymorphism in endoplasmic reticulum antigen processing peptidases (ERAAP) that trim MHC class I ligands is associated with autoimmune diseases, but the mechanism is unknown. We analyzed the effect of Eraap deficiency in the non-obese diabetic (NOD) mouse model of type 1 diabetes. *Eraap*^*-/-*^ NOD mice displayed reduced and delayed diabetes, harbored splenic effector T cells unable to transfer diabetes, and exhibited a strong shift from effector to central memory T cells in attenuated islet infiltrates. Eraap deficiency increased presentation of the immunodominant epitope insulin B_15-23_ by beta cells but at the same time provided complete protection from diabetes to chimeras reconstituted with bone marrow encoding a CD8^+^ T cell recognizing this epitope. These results underline the strong impact of self-antigen presentation to CD8^+^ T cells in diabetes. At the same time, they highlight the complex consequences of interfering with MHC-I antigen presentation in autoimmunity and advise caution in therapeutic modulation of ERAP activity in this context.

## Introduction

Autoimmune diseases often implicate recognition of self-antigen peptides presented by MHC class I and class II molecules to CD8^+^ and CD4^+^ T lymphocytes, respectively [1,2]. The specific peptide binding preferences of individual MHC molecules, based on their genetic polymorphism, have long been known to affect the likelihood of developing autoimmune conditions, resulting in firmly established associations of defined MHC-I and MHC-II allomorphs with diseases such as ankylosing spondylitis or type 1 diabetes (T1D), respectively [3]. More recently, polymorphism in genes encoding the endoplasmic reticulum aminopeptidases ERAP1 and ERAP2 have also been found to be associated with the risk of several autoimmune diseases [4]. These include ankylosing spondylitis, psoriasis, Behçet’s disease, and birdshot chorioretinopathy, associated with ERAP1 and ERAP2 variants, and juvenile idiopathic arthritis associated with ERAP2 only [5]. T1D has also been suggested to be associated with ERAP polymorphism although this is controversial [6]. However, the mechanism underlying these associations remains unknown.

Human ERAP1 and ERAP2 and mouse ERAAP play a key role in MHC-I antigen presentation by aminoterminal “trimming” of epitope precursors. Trimming can produce peptides of optimal length for MHC-I binding but also destroy potential epitopes [7]. As a result, ERA(A)P activity has a substantial impact on the immunopeptidome presented by antigen presenting cells (APCs). This is illustrated by the strong CD8^+^ T cell response triggered by ERAAP-deficient cells injected in wild type mice [8], and by the antigenicity of tumor cells treated with ERAP inhibitors, an effect with potential for therapeutic application in cancer [9,10]. However, while global effects on the immunopeptidome can be of obvious therapeutic interest in cancer, they are less likely to explain how ERAP variants can promote or prevent autoimmune CD8^+^ T cell responses. Conversely, in autoimmunity, a possible scenario is that variable ERAP trimming has a massive impact on presentation of one or a small number of key self-epitopes presented by an MHC-I molecule itself associated with disease. Efficient trimming by a disease-associated ERAP1 variant of a melanocyte epitope presented by HLA-C*06:02, an allomorph predisposing for psoriasis, could correspond to this scenario [11]. Similarly, we found that ERAP1-dependent presentation of a single model epitope can have a dramatic impact on the entire array of MHC-I molecules displayed by an APC [12].

To decipher mechanisms underlying *ERAP*-linked predisposition to autoimmune diseases, animal models can be useful. Knockout of *Eraap* in a mouse strain not predisposed for autoimmunity has been reported to induce hallmark features of ankylosing spondylitis such as ankylosis and spinal inflammation [13]; epitopes processed by ERAAP in this model were not reported. We investigated the effect of *Eraap* deficiency in non-obese diabetic (NOD) mice, a model of spontaneous autoimmunity in which a key CD8^+^ T cell self-epitope is known [14,15]. We report the unexpected finding that ERAAP deficiency enhances presentation of this key epitope by beta cells but protects against transfer of diabetes by cognate CD8^+^ T cells as well as from spontaneous disease. These results underline the complexity of the role of ERA(A)P trimming in autoimmunity and advise caution in modulating enzyme activity in this context.

## Materials and Methods

### Generation of Eraap^-/-^ NOD mice

*Eraap*^*-/-*^ mice were produced by injecting NOD zygotes with CRISPR/Cas9 components and transfer of the injected embryos in pseudopregnant females. Two SgRNA guides (SgRNA1 UGAAUCGGGGUCAUAUACUC; SgRNA2 GGAAGUCGCAUAUUAUUCCA) targeting exon 2 of *Eraap* were injected simultaneously. Genome-modified mice were identified by PCR amplification and sequencing of exon 2. Genomic DNA was extracted from mouse tails with the Kapa Express Extract kit (Sigma Aldrich) and amplified using the primers: Forward: 5’AAGAGAGATCCCAAATGGCAAACAGAA-3’; and Reverse: 5’GCCTTAGATATTTGCAGGTGGTGACTA-3’. The PCR product was sequenced by Eurofins Genomics. Mutations at the genomic *Eraap* locus were analyzed by MacVector (version 18.2.5) software. A homozygous founder was selected that exhibited a 49 base pair deletion in exon 2 at the *Eraap* SgRNA2 DNA strand break site. This deletion resulted in a frame shift at amino acid 41 and the creation of an amber mutation 15 bp downstream. Germline transmission of *Eraap* mutation was identified by PCR on offspring DNA. Ethical approval for animal experimentation was accorded by the *Ministère de l’enseignement, de la recherche et de l’innovation* on January 29, 2024, for a duration of 5 years under the number APAFIS #46924-2023102612559010.

### Diabetes measurement

The onset of diabetes was defined as two positive urine glucose tests with a dipstick test, confirmed by a glycemia >250 mg/dL using Glucotest (Roche) and performed in a blinded fashion. Glucosuria was monitored three times per week. Following an initial positive glycosuria test, glycemia was monitored twice weekly until the age of 30 weeks or until the humane end point of glycemia > 600 mg/dl was reached.

### Insulitis scoring

Insulitis was assessed in 16-week-old female NOD-*Eraap*^*-/-*^ mice and age-matched controls with three mice per group. Pancreata were fixed in 4% formalin for 24h and subsequently paraffin-embedded, sectioned at 4 μm thickness, and stained with hematoxylin-eosin. Sections were blindly evaluated for insulitis severity using the following scale: 0 - normal, 1 - minor peri-insulitis, 2 - extensive peri-insulitis affecting more than 50%, 3 - moderate intra-insulitis with less than 50% mononuclear cell infiltration, and 4 - severe insulitis characterized by over 50% mononuclear cell presence and/or disrupted islet architecture. A total of 25 and 28 islets from three mice per genotype were analyzed, respectively.

### Adoptive transfer of effector T cells

Spleens of *Eraap*^*-/-*^ and age-matched control mice were first depleted of non-CD3^+^ T cells using the MojoSort™ Mouse CD3 T Cell Isolation Kit (Biolegend). The remaining cells were stained for CD25, CD62L and CD3, and CD3^+^CD62L^-^CD25^-^ cells were sorted using a FACSAria cell sorter (BD Biosciences). Finally, 5x10^5^ cells in 100 μL PBS were administered into the retro-orbital plexus of female *Rag1*^*-/-*^ NOD mice aged seven weeks under gaseous anesthesia. Glycemia was monitored three times per week until the endpoint.

### Bone marrow chimeras

*Eraap*^*-/-*^ NOD mice aged 10 weeks and matched control females were treated with 30 mg/kg Busulfan for a myeloablative conditioning over two days before retro-orbital injection of 5 x 10^6^ G9C8 bone marrow cells in 100 μL PBS. Bone marrow was harvested from the hind legs of TCR-transgenic G9C8 mice (NOD.Cg-Tcratm1MjoTg(TcrbG9C8)2Fsw/J(NOD-TgG9C8) aged 11-12 weeks. Glycosuria and glycemia were monitored as stated above.

### Flow cytometry

Cells from pancreatic islets were isolated as previously described (Bertocci et al., 2025) from *Eraap*^*-/-*^ NOD mice aged 12 weeks and matched controls. Staining for flow cytometric analysis was performed in 100 μl of Stain Buffer (BD Bioscience) with the antibodies reported in table S1. The cells were washed twice with 2 ml Stain buffer, resuspended in PBS/2% FCS and analyzed using a SonyID7000 spectral analyzer. FlowJo software (version 10.8.2) was used for analysis.

## Results

To interrogate the effect of Eraap deficiency on autoimmune diabetes in the spontaneous NOD model, we undertook a CRISPR-Cas9-mediated knockout in NOD zygotes; this avoided the requirement of a lengthy backcross and potential effects of carried-over genes from a non-autoimmune background. Of 5 founder mice obtained, we selected a mouse with a homozygous frame shift mutation in exon 2 of the *Eraap* gene, resulting in a predicted truncation of any protein product at amino acid 46 (Fig. 1A).

**Figure 1.**
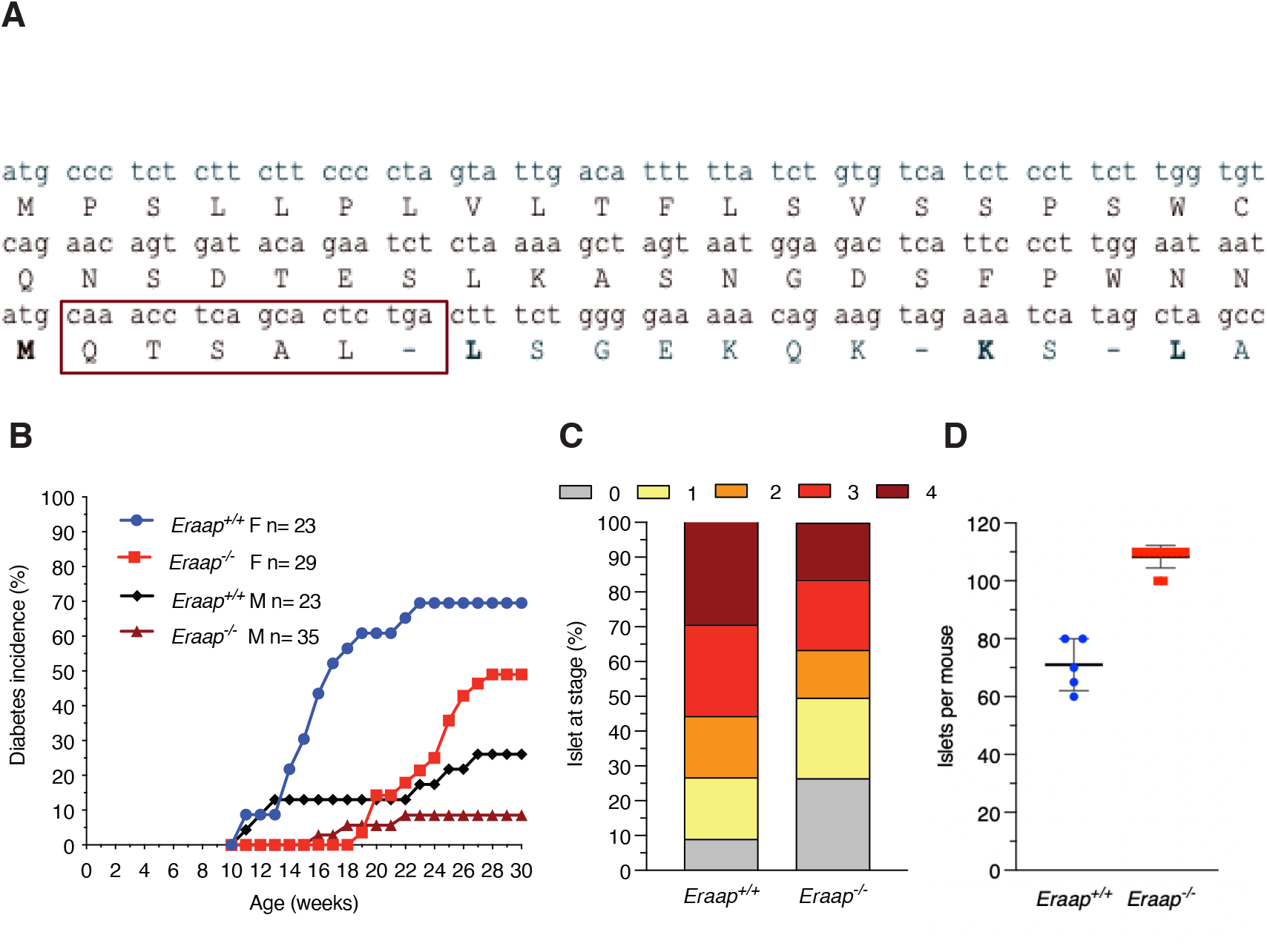
Eraap deficiency confers protection from diabetes to NOD mice. **(A)** Exon 2 in *Eraap*^*-/-*^mice. Amino acids and stop codon downstream of the frame shift are boxed. **(B)** Diabetes incidence in *Eraap*^*-/-*^ and wild-type cohorts. p<0.0002 at 26 weeks by Kaplan-Meier, log rank (Mantel-Cox) test. **(C)** Islets from 16-week-old female *Eraap*^*+/+*^ and *Eraap*^*−/−*^ mice (n=3) were blindly evaluated for insulitis. Scale: 0 = no infiltrate, 1 = peri-islet infiltrate, 2 = extensive (>50%) peri-islet infiltrate, 3 = intra-islet infiltrate, and 4 = extensive/destructive intra-islet infiltrate (>50%). p=0.005 by Chi-square test. **(D)** Hand-picked islets from 12-week-old female *Eraap*^*−/−*^ (n=5) and aged-matched *Eraap*^*+/+*^ (n= 12) were enumerated.

To assess the effect of Eraap deficiency on diabetes, we compared the diabetes in female and male homozygous *Eraap*^*-/-*^ mice with wild-type (WT) controls. While manifest diabetes started appearing at 11 weeks and reached an incidence of 70% in WT female mice, disease development was delayed by 8 weeks and reached only 49% in female mutant mice.

Similarly, diabetes was delayed (11 vs. 16 weeks) and reached a lower plateau (8.6 vs. 26%) in male *Eraap*^*-/-*^ mice (Fig. 1B). Scoring of islet infiltration showed an attenuation of insulitis in 16-week-old *Eraap*^*-/-*^ mice, with 16 vs. 30% in WT mice strongly infiltrated and 26% vs. 8% devoid of infiltration (Fig. 1C). Consistent with this, islets were more numerous in mutant mice (110 vs. 70 islets per mouse) (Fig. 1D). Thus, Eraap deficiency provides partial protection from diabetes and insulitis in NOD mice.

Protection from diabetes can be due to absence of diabetogenic effectors and/or increased activity of protective regulatory T cells. Diabetogenic effectors can be purified as CD3^+^CD25^-^ CD62L^-^ cells from NOD spleens [16]. In 2 independent experiments with a total of 10 donors of splenocytes per genotype, splenic effectors from WT mice induced diabetes in 100% of *Rag1*^*-/-*^ NOD recipients, while splenic effector T cells from *Eraap*^*-/-*^ donors were completely unable to do so (Fig. 2A).

**Figure 2.**
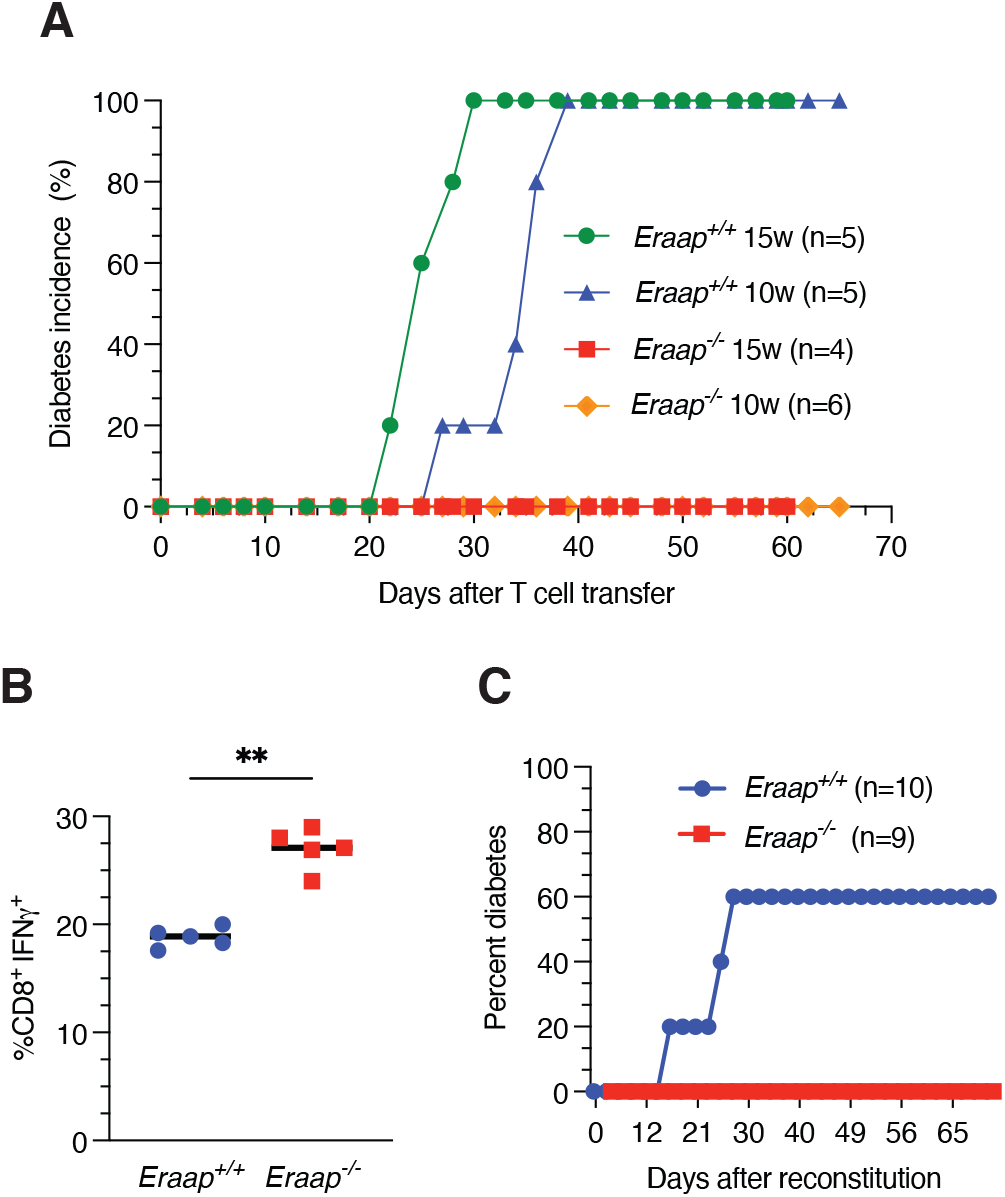
ERAAP deficiency protects against diabetes transfer by poly- and monoclonal T-cell populations. **(A)** Adoptive transfer of splenic effector T cells isolated from *Eraap*^*+/+*^ or *Eraap*^*−/−*^ donors (aged 10 or 15 weeks) to *Rag1*^*-/-*^ NOD recipients. Statistical analysis by Kaplan-Meier, log rank (Mantel-Cox test), p<0.0001. **(B)** IFN-γ secretion by G9C8 CD8^+^ T cells stimulated with live islets from 16-week-old female *Eraap*^*+/+*^ and *Eraap*^*−/−*^ NOD mice was assessed by flow cytometry. **(C)** *Eraap*^*-/-*^ and *Eraap*^*+/+*^ mice were reconstituted with bone marrow cells from G9C8 TCR-transgenic mice and monitored for diabetes onset. The results shown are cumulated from two independent experiments.

Autoimmune diabetes results from killing of NOD beta cells by CD8^+^ T cells, in which T cells recognizing the insulin B_15-23_ epitope play a key immunodominant role at least in early insulitis [17–19]. We tested recognition of WT and *Eraap*^*-/-*^ beta cells by transgenic G9C8 cells expressing a T cell receptor recognizing this epitope [17]. *Eraap*^*-/-*^ beta cells stimulated IFN-γ release by G9C8 cells more efficiently than WT beta cells (Fig. 2B), confirming our previous results obtained with *Eraap*^*-/-*^ NOD beta cells obtained through crossing with *Eraap*^*-/-*^ C57BL/6 mice [16]. This suggests that Eraap tends to destroy the B_15-23_ epitope, reducing its presentation.

Next, we asked how increased presentation of B_15-23_ affects diabetes development in NOD mice. WT NOD chimeras grafted with G9C8 bone marrow developed disease by four weeks after reconstitution, consistent with the pathogenicity of B_15-23_-specific T cells [17]. In contrast, *Eraap*^*-/-*^ recipients of G9C8 bone marrow were surprisingly completely protected from disease (Fig. 2C).

Considering that many *Eraap*^*-/-*^ mice eventually developed disease, contrasting with complete protection upon adoptive splenocyte transfer and G9C8 chimera production, we finally examined islet infiltrates and pancreatic lymph node (PLN) cells to search for diabetogenic effectors. Flow cytometric analysis (Suppl. Fig. 1) did not reveal any significant changes among PLN lymphocytes (Suppl. Fig. 2). In contrast, *Eraap*^*-/-*^ islets contained reduced numbers of CD45^+^ immune cells (Fig. 3A) as well as of B, CD4^+^ and CD8^+^ T lymphocytes (Fig. 3C, E, M) while the percentage of each of these populations among total CD45^+^ cells was unchanged (Fig. 3B, D). Both the percentage and the number of naïve (CD44^-^CD62L^+^) CD4^+^ and CD8^+^ T cells were increased in *Eraap*^*-/-*^ islets (Fig. 3F, G, N, O). Moreover, both among CD4^+^ and CD8^+^ T lymphocytes, the ratio between effector memory (CD44^+^CD62L^-^) and central memory (CD44^+^CD62L^+^) cells was strongly altered in *Eraap*^*-/-*^ islets, such that the percentage and the number of the former were reduced while the opposite was the case for the latter (Fig. 3H, I, J for CD4^+^; 3P, Q, R for CD8^+^). Interestingly, infiltrates in *Eraap*^*-/-*^ islets also comprised an increased percentage of CD4^+^CD62L^+^CD25^+^ presumable regulatory T cells (Fig. 3K).

**Figure 3.**
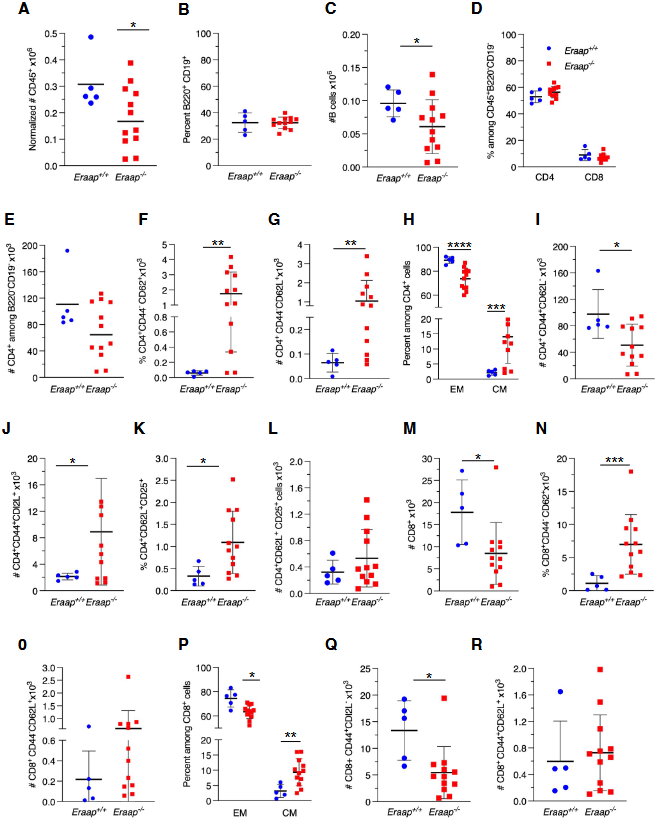
ERAAP deficiency reduces and alters immune cell islet infiltration. Single-cell suspensions of islet-infiltrating cells from 12-week-old female *Eraap*^*−/−*^ mice and age-matched control *Eraap*^*+/+*^ mice were analyzed by flow cytometry. Cell numbers were normalized to 10^6^ cells acquired for each mouse. **(A)** Number of CD45^+^ cells. **(B)** Percent among CD45^+^ cells and **(C)** number of B220^+^CD19^+^ B cells. Percent among non-B cells **(D)** and number of CD4^+^ **(E)** and CD8^+^ **(J)** cells. Percent among CD4^+^ cells **(F)** and number of CD4^+^ effector memory **(G)** and central memory **(H)** T cells. **(I)** Percent of CD4^+^ T regulatory cell among CD45^+^ cells. Percent among CD8^+^ cells **(K)** and number of effector **(L)** and central **(M)** memory CD8^+^ T cells. Each symbol represents one mouse. Data (mean ± SD) are from two independent experiments. Statistical analysis was performed with unpaired, two-tailed t-test with Welch correction. *p <0.05,**p <0.01, ***p <0.001, ****p<0.0001.

## Discussion

The genetic association of ERAP1/2 with autoimmune diseases has raised interest in using the enzymes as targets for therapeutic or prophylactic intervention [20]. This study was undertaken to explore potential effects of such approaches. Our choice of the NOD model was motivated by the spontaneous nature and the important role of CD8^+^ T cells in the disease, but also by the existence of an immunodominant self-epitope with a clear role in disease initiation [14]. Next to this, the potential association between ERAP polymorphism and T1D was a secondary and minor reason for the choice [6]. Our results highlight the complexity of interfering with MHC-I antigen presentation in autoimmunity.

The global result of Eraap deficiency was partial protection from diabetes that was strongly delayed and reduced in incidence, underlining the key role of CD8^+^ T cells recognizing MHC-I-presented self-epitopes in initiation and amplification of the autoreactive beta cell attack [19]. Relative protection from disease correlated with an absence of diabetogenic potential of splenocytes and an increase in islet-infiltrating regulatory CD4^+^ T cells. Islet infiltrating *Eraap*^*-/-*^ T cells also displayed a remarkable shift from effector memory to central memory phenotype, associated with a global decline in infiltration and an increased number and percentage of naïve T cells. While we do not know the mechanism underlying this shift, dominance of effector memory over central memory cells has been shown in other models to be associated with a stronger autoaggressive profile as evidenced by higher inflammatory cytokine production and proliferation [21–24].

Our findings suggest that altered presentation of the immunodominant insulin epitope B_15-23_ may play an important role in relative protection from disease. Complete protection from disease suggests that chimeras lacked the early CD8^+^ T cell response likely initiating T cell autoimmunity, explaining the delayed disease onset presumably driven by other T cell specificities. The reason for protection of chimeras remains to be determined. Activated G9C8 cells can transfer diabetes to NOD.scid mice [17]. *Eraap* is expressed by both beta cells and thymic antigen presenting cells which also express insulin [25] at relatively high levels (www.bgee.org./gene/ENSMUSG00000021583). We speculate that enhanced presentation of insulin B_15-23_ by thymic APCs may promote deletion or tolerization of G9C8 cells, blunting their autoreactive potential. Whatever the mechanism of protection, our results show that i) interfering with MHC class I epitope trimming can have a substantial impact in a T cell-mediated autoimmune disease, and ii) the nature of this impact can be complex and difficult to deduce from the effect of Eraap on presentation of immunodominant self-epitopes.

## CRediT authorship contribution statement

B. Bertocci: Investigation, formal analysis, visualization, writing - review & editing; E. Waeckel-Énée: Investigation, formal analysis; N. Keelan: Investigation; S. You: Resources, writing - review & editing; P. David: Investigation; P. van Endert: Conceptualization, funding acquisition, project administration, supervision, writing - original draft & editing.

## Acknowledgements

the authors are grateful to Nicolas Kuperwasser (Gene editing and targeting platform, Institut Necker Enfants Malades) for the design of *Eraap-*targeting guide RNAs.

## Data availability

data will be made available upon request.

