## Supplementary material for "Interference with MHC class I epitope trimming provides paradoxical protection from autoimmune diabetes": Suppl. table 1 and suppl. figures 1 and 2

**Supplementary Figure 1**: *Gating strategy for flow cytometry analysis of lymphocytes in islets and lymph nodes*. (**A, B**) Singlet CD45$^{+}$ cells were first separated according to CD19 and B220 expression. B220-negative cells were then stained with antibodies to CD4 and CD8 and further divided in effector memory (EM), central memory (CM), naive and regulatory T cells by staining for CD44, CD62L and CD25.

**Supplementary Figure 2**: (**A-R**) the percentage and the number of B cell and T cell populations in pancreas-draining lymph nodes of 12-week-old female *Eraap*$^{+/+}$ and *Eraap*$^{-/-}$ mice was analyzed exactly as shown for islet-infiltrating lymphocytes in Fig. 3.

**Supplementary table 1**: Antibodies used

| Antibody | Fluorochrome | Clone | Dilution | | Catalogue number |
|---|---|---|---|---|---|
| CD45 | Percp | 30F11 | 1/130 | Biolegend | 103130 |
| CD3 | APC | 17A2 | 1/100 | ThermoFischer Scientific | 17- -0032-80 |
| B220 | PEeFluor610 | RA3-6B2 | 1/130 | ThermoFischer Scientific | 61-0452-82 |
| CD19 | APC-eFluor780 | eBio1D3 | 1/130 | ThermoFischer Scientific | 47-0193-82 |
| GL7 | eFluor450 | GL7 | 1/130 | ThermoFischer Scientific | 48-5902-82 |
| CD95 | PE-Cy7 | Jo2 | 1/130 | BD Bioscience | 557653 |
| CD1d | SB600 | 1B1 | 1/130 | ThermoFischer Scientific | 63-0011-82 |
| CD5 | Percp-Cy5.5 | 53-7.3 | 1/130 | Biolegend | 100623 |
| CD86 | PE-Cy5 | GL1 | 1/130 | ThermoFischer Scientific | 15-0862-81 |
| CD4 | BV241 | GK1.5 | 1/130 | Biolegend | 100443 |
| CD8a | APC | 53-6.7 | 1/130 | Sony | 1103560 |
| CD62L | BV785 | MEL-14 | 1/130 | Biolegend | 104440 |
| CD62L | Biotin | MEL-14 | 1/100 | ThermoFischer Scientific | 13-0621-82 |
| CD25 | PE | PC61.5 | 1/130 | | 12-0251-81 |
| CD11c | BV711 | N418 | 1/130 | Biolegend | 517310 |
| CD11b | BV510 | M1-70 | 1/130 | Biolegend | 101263 |
| MHCII I-A[k] | FITC | 10-306 | 1/130 | BD Bioscience | 562014 |
| F4/80 | AF700 | BM8 | 1/60 | ThermoFischer Scientific | 56-4801-80 |
| CD44 | BV650 | IM7 | 1/130 | BD Bioscience | 740455 |
| Zombie NIR$^{TM}$ Fixable Viability Dye | | | 1/100 | Biolegend | 423105 |
| Sytox blue | | | 1/400 | ThermoFischer Scientific | S34857 |
| Streptavidin | APC-eFluor-780 | | 1/200 | ThermoFischer Scientific | 47-4317-82 |

**A**

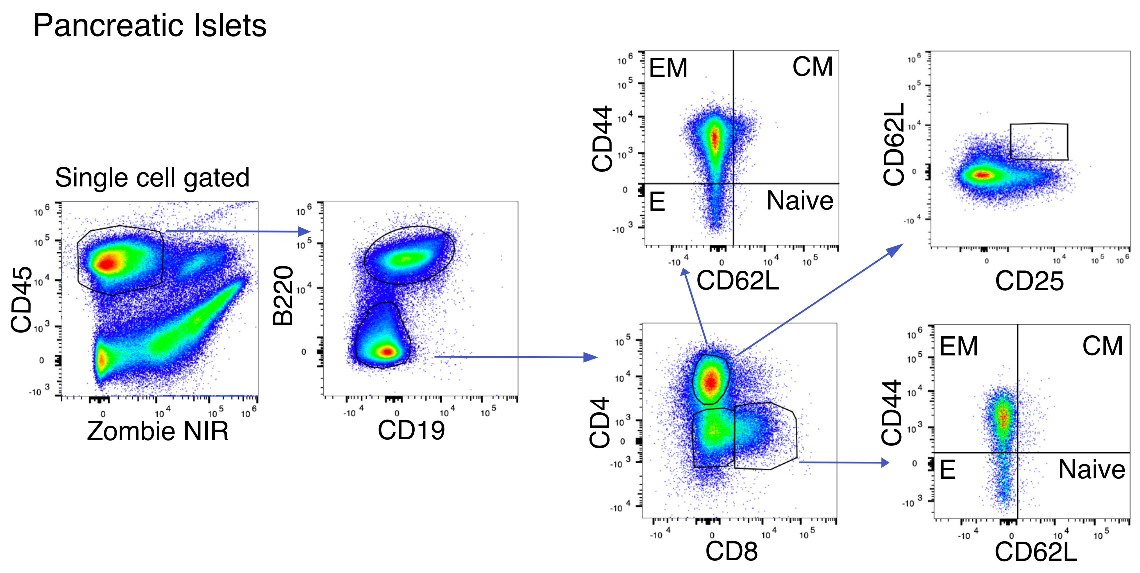
Pancreatic Islets
Single cell gated
CD45
Zombie NIR
B220
CD19
CD4
CD8
CD44
CD62L
EM
CM
E
Naive
CD62L
CD25
CD44
CD62L
EM
CM
E
Naive

**B**

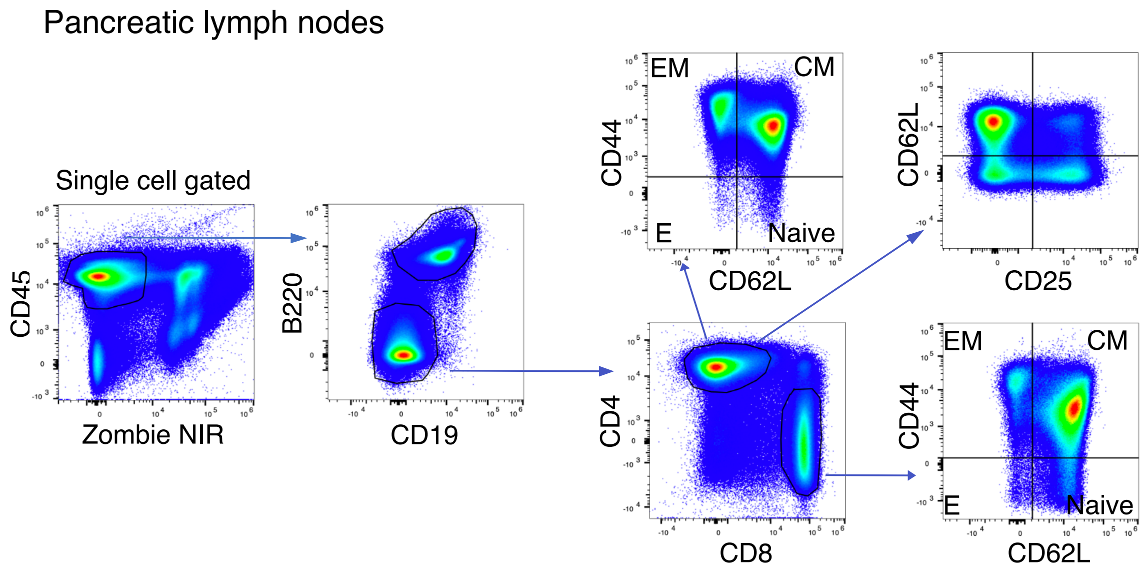
Pancreatic lymph nodes
Single cell gated
CD45
Zombie NIR
B220
CD19
CD4
CD8
CD44
CD62L
EM
CM
E
Naive
CD62L
CD25
CD44
CD62L
EM
CM
E
Naive

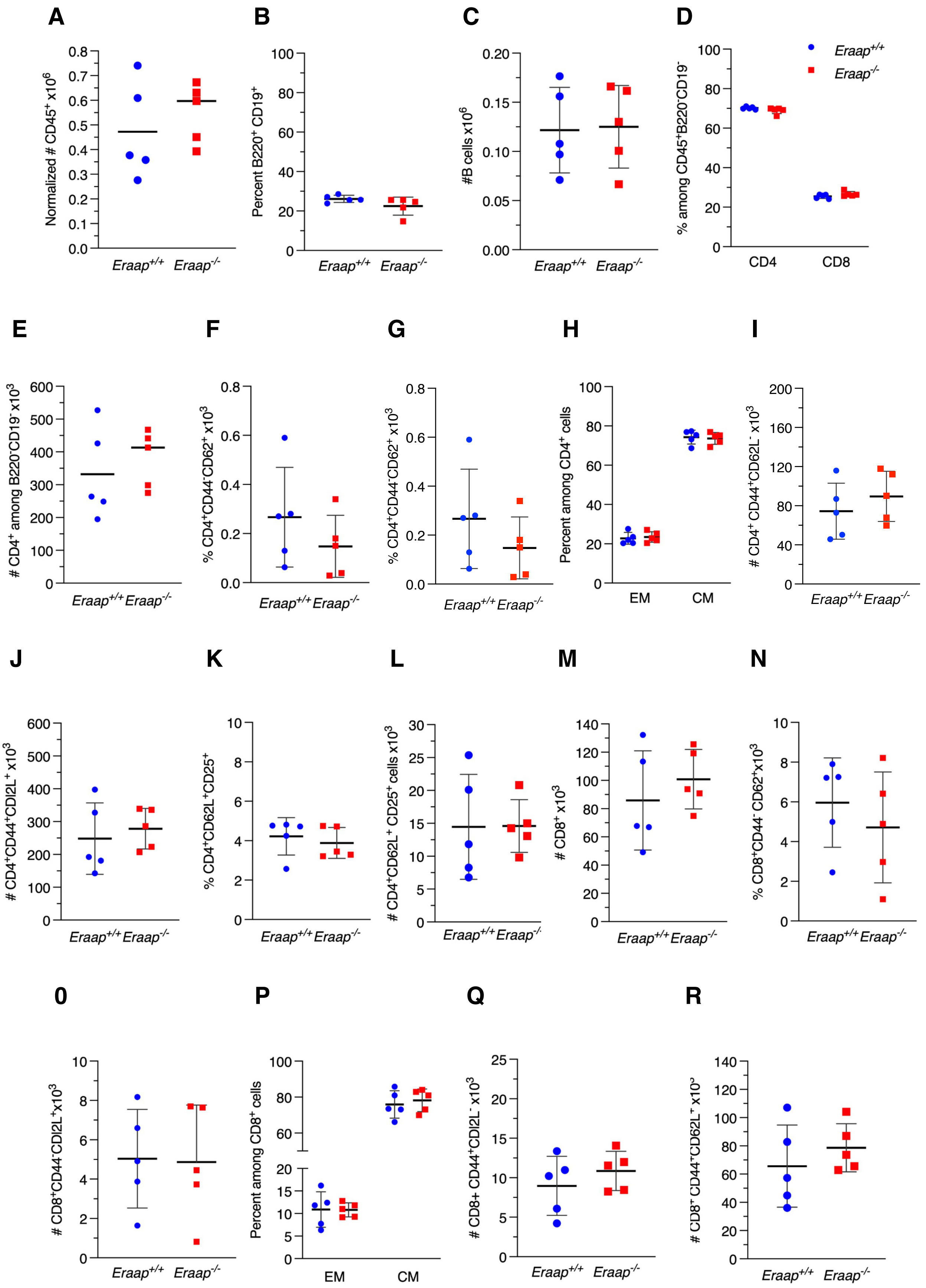
A
Normalized # CD45+ x10^6
Eraap+/+ Eraap-/-
B
Percent B220+ CD19+
Eraap+/+ Eraap-/-
C
#B cells x10^6
Eraap+/+ Eraap-/-
D
% among CD45+B220-CD19-
Eraap+/+
Eraap-/-
CD4 CD8
E
### CD4+ among B220-CD19- x10^3
Eraap+/+ Eraap-/-
F
% CD4+CD44-CD62+ x10^3
Eraap+/+ Eraap-/-
G
% CD4+CD44-CD62+ x10^3
Eraap+/+ Eraap-/-
H
Percent among CD4+ cells
EM CM
I
# CD4+ CD44+CD62L- x10^3
Eraap+/+ Eraap-/-
J
### CD4+CD44+CDl2L+ x10^3
Eraap+/+ Eraap-/-
K
% CD4+CD62L+CD25+
Eraap+/+ Eraap-/-
L
### CD4+CD62L+ CD25+ cells x10^3
Eraap+/+ Eraap-/-
M
# CD8+ x10^3
Eraap+/+ Eraap-/-
N
% CD8+CD44- CD62+x10^3
Eraap+/+ Eraap-/-
0
### CD8+CD44-CDl2L+x10^3
Eraap+/+ Eraap-/-
P
Percent among CD8+ cells
EM CM
Q
### CD8+ CD44+CDl2L- x10^3
Eraap+/+ Eraap-/-
R
# CD8+ CD44+CD62L+ x10^3
Eraap+/+ Eraap-/-
